# Maintenance of viral diversity through influenza transmission bottlenecks: a within-host branching-process model

**DOI:** 10.64898/2026.06.24.734279

**Authors:** Wenjing Zhang, Leif Ellingson, Lisa Bono

## Abstract

Viral populations can experience a dramatic reduction in population size and *genetic diversity* during transmission between donor and recipient hosts. *Transmission bottlenecks* can therefore decrease the evolutionary potential of viral populations, slowing *adaptation* by increasing the strength of *genetic drift* and decreasing the strength of selection. Recent barcoded influenza experiments in guinea pigs showed that recipient animals receive a diverse viral inoculum but lose most of that diversity within one to two days. The resulting bottleneck therefore arises not at physical transfer but during early viral growth in the recipient. We develop a branching-process framework to quantify how much of this loss follows from stochasticity in early growth alone. Each transmitted lineage is treated as an independent stochastic process governed by measurable viral parameters. We recover these parameters from viral growth rates estimated from observed viral load. A residual filter for each animal then captures any additional loss imposed by the recipient host. Applied to twenty-four recipient animals, the model reveals two distinct groups. For roughly half of the animals, the stochastic extinction during early growth already accounts for the observed loss. For the remaining animals, the additional host filter is severe. Only about one in a hundred free *virions* pass through. This decomposition offers a quantitative entry point for future work on immune contributions to transmission bottlenecks.

## 1 Introduction

Viral diversity can decline at three stages of transmission: within the donor host, in the environment between hosts, and within the recipient host. Recent experimental work suggests that for some viruses the primary driver of the bottleneck is the third stage, the earliest period of within-host establishment in the recipient [15]. This raises a broader mechanistic question. How much of the observed loss follows from chance alone, and when must filtering by the recipient host be invoked?

### 1.1 Background

Genetic diversity is essential for viral *evolution*. During infection, a viral population can balloon to immense sizes. With every progeny *virion* produced during this population expansion, there is the potential for new genetic variation to arise. Viruses, especially RNA viruses, have high *mutation* rates and readily exchange genetic material [9]. Thus, not only do the populations increase in size but also in genetic diversity. This novel variation allows the viral population to explore sequence space, potentially finding new *genotypes* that are better adapted to their environment [39]. The founding population of each new infection determines the standing genetic variation on which selection can act and sets an upper bound on the adaptive potential available for immune escape, *antigenic variation*, and drug resistance [22, 39]. These large populations are subject to selection that can prune less fit variants and increase the overall *fitness* of the population in a deterministic fashion.

However, when the viral population is transmitted from a donor host to a recipient host, its size is sharply reduced. Only a small fraction of the donor population successfully initiates infection in the recipient [39], and the genetic composition of these founders differs from that of the donor in the number and frequency of variants [38]. Such a loss of diversity is a *genetic bottleneck*, the classical population-genetics phenomenon in which a sharp reduction in population size compresses genetic diversity. A *transmission bottleneck* is the special case in which the loss occurs during transfer of an infection from one host to the next. Some infections are founded by just a few or even a single infectious viral particle, e.g. poliovirus [34], hepatitis C virus [37], and several plant viruses [4, 21]. With its small size, this founding population of viruses is more vulnerable to the stochastic effects of genetic drift, which prunes variants at random, regardless of their fitness effects, and decreases the overall fitness of the viral population. This means that not only are deleterious and neutral variation purged, but beneficial genotypes can also be lost during these genetic bottlenecks, which can slow adaptation, especially for viruses emerging in novel environments [22].

Diversity can be lost at more than one point along the transmission process, and different models have emphasized different points. A recent model attributes the bottleneck primarily to the physical airborne transfer between donor and recipient, particularly for respiratory viruses [31]. This focus on the transfer step leaves open a complementary contribution from within-host dynamics. Loss can also occur within the recipient host, through immune or physical filtering in the airway (the *recipient-side filter*) or through *stochastic dynamics* during the earliest cycles of within-host replication, with genetic drift acting throughout to remove variants at random regardless of fitness. A further source of loss is that some transmitted particles are non-infectious to begin with, such as virions carrying lethal mutations or defective interfering particles [39].

Each of these losses can also fall on variants at random or according to their fitness, and determining the relative roles of drift and selection has proven difficult, in part because the two forces appear to dominate at different scales. Transmission bottlenecks dramatically reduce population size and diversity, conditions under which drift would be expected to dominate. Consistent with this expectation, within individual hosts genetic drift and *purifying selection* appear to dominate during a single infection [23, 13, 8, 32], as also observed for SARS-CoV-2 [33, 12, 6, 19]. Yet at the scale of the whole human population, selection is clearly detectable: seasonal influenza shows a signature of *positive selection*, with beneficial mutations sweeping the global population and driving antigenic adaptation [10, 20, 11].

Few studies, however, are designed to separate the loss caused by drift from that caused by selection. A recent elegant experiment was designed to do exactly this, using genetically identical influenza A genotypes that differ only by neutral barcodes to isolate drift and minimize selection. Guinea pigs were inoculated with these neutrally barcoded viruses and the diversity of the viral populations was tracked through infection and transmission. Genetic diversity remained high in the donor host and for one to two days post-infection in the recipient, suggesting that the genetic bottleneck arises from host factors within the recipient [15]. Here, we use a branching-process framework to recapitulate transmission of the neutrally barcoded population between guinea pigs and to quantify how much of this lineage loss is attributable to stochastic growth dynamics alone.

### 1.2 Empirical context

Holmes *et al*. used an Influenza A/Panama/2007/99 (H3N2) virus (Pan/99) library carrying up to 4,096 neutral nucleotide barcodes. They inoculated a donor guinea pig and allowed transmission to a recipient through both direct contact and aerosol, sampling barcode diversity every day. Three key findings emerged. First, inoculated guinea pigs maintain high barcode diversity throughout infection, indicating that within-host dynamics in the donor do not impose a severe bottleneck. Second, recipients of both direct-contact and aerosol transmission begin with surprisingly high diversity on their first day of infection, confirmed by plaque assay, implying that many viral genotypes successfully cross the environmental barrier. Third, this diversity collapses within one to two days, with Bray–Curtis dissimilarity between successive days decaying rapidly thereafter.

These observations led Holmes *et al*. to conclude that the early viral expansion in the recipient, rather than the physical transfer event itself, is a primary driver of the influenza A transmission bottleneck. They speculated that target-cell limitation, superinfection exclusion, spatial separation, or early immune effectors may contribute to the diversity loss.

They further proposed that this loose first stage of transmission could provide a previously unrecognized opportunity for selection, because the incoming population is still antigenically diverse when it encounters recipient immune defenses. The magnitude of this selective opportunity, however, remains to be quantified, and doing so requires a mechanistic model that can predict how much contraction should occur from stochastic dynamics alone, explain why contraction severity varies so widely across animals, and partition the observed loss into its contributing causes.

The finding that viral expansion after transfer is a primary driver of the bottleneck [15] is qualitative, however, and has not yet been developed into a quantitative theory. Four questions remain open. First, how much of the observed contraction can be attributed to stochastic growth and clearance alone, without invoking any immune or physical filtering? Second, how does bottleneck strength depend on the biological parameters governing replication and clearance? Third, why does the observed contraction vary by two orders of magnitude across recipient animals? Fourth, for animals whose diversity loss exceeds what growth-only dynamics predict, how strong must the additional recipient-side filter be?

### 1.3 Modelling context

Answering these questions requires a stochastic framework in which lineage extinction probabilities depend explicitly on viral life-history parameters. Such a framework has been developed over the past two decades by Wahl and collaborators in the context of fixation-probability theory. Wahl and Gerrish [35, 36] showed that periodic population bottlenecks profoundly alter fixation probabilities, with the effect depending on the timing and severity of the bottleneck relative to the growth phase. Hubbarde, Wild, and Wahl [17] introduced the burst-death model, in which generation times are exponentially distributed and each individual produces a fixed burst of offspring, and derived the probability generating function (pgf) governing lineage dynamics. They showed that fixation probabilities under this life history differ substantially from classical predictions, and Hubbarde and Wahl [16] subsequently used the framework to determine optimal bottleneck ratios for experimental evolution. Patwa and Wahl [25] extended the model to the attachment-lysis framework for lytic viruses, incorporating an explicit attachment rate, a clearance rate, a fixed eclipse period, and an integer burst size, and solved the resulting delay differential equation for the pgf by the method of stepwise integration [14]. Further extensions addressed host-cell dynamics [26], the dependence of fixation on which life-history trait is affected by a mutation [1], and the broader theoretical landscape of fixation probabilities [27]. We adapt their pgf framework to the complementary problem of tracking existing barcode lineages through the post-transfer establishment phase, rather than predicting the fate of new mutations over the full infection cycle.

### 1.4 Contributions and organization

We adapt this framework to obtain the missing quantitative parts. A stability-constrained inversion connects the estimated growth rate *λ* to the mechanistic infection rate *A*, bridging the titer and barcode diversity data collected by Holmes *et al*. A per-animal residual filter 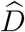 then partitions each recipient’s diversity loss into a stochastic-growth component and a host-imposed component. The composite map *λ* ↦ 1 − *q*, where *q* is the ultimate lineage extinction probability, spans a 95-fold range across the observed *λ* distribution, accounting for the wide between-animal variation without requiring per-animal differences in immune status.

Section 2 develops the branching-process model and the inversion *λ* ↦ *A*. Section 3 quantifies the empirical lineage contraction across the 24 recipient animals of [15]. Section 4 introduces the recipient-side filter and reports the per-animal estimates 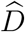. Section 5 discusses implications for virus-immune modeling and future extensions. Supporting derivations and per-animal tables appear in Appendices A and B.

## 2 Mathematical Modelling

### 2.1 Branching-process modeling framework

At early stages of viral expansion, viral titers remain low relative to the large pool of susceptible epithelial cells, so most infected cells are likely initiated by single virions and coinfection is rare. Over the short timescales considered here, density-dependent and immune feedbacks are also expected to be weak. We therefore assume that each barcode lineage can be approximated by an independent stochastic branching process, as in attachment–lysis models for lytic viruses [26].

#### Branching-process formulation

Let *N* (*t*) denote the number of free virions in a lineage founded by a single virion at time *t* = 0, and let *G*(*x, t*) = E[*x*^*N*(*t*)^] be its pgf. A free virion infects a target cell at rate *A* or is removed at rate *C*. Following infection, a fixed latent period *T*_*L*_ elapses before the infected cell releases a burst of *B* free virions. Under the assumption that descendant lineages evolve independently, *G* satisfies [25]

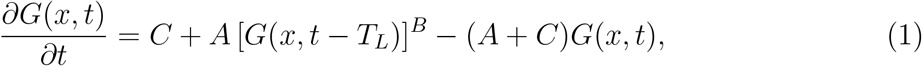

with initial conditions *G*(*x*, 0) = *x, G*(*x, t*) = 1 for −*T*_*L*_ ≤ *t* < 0.

Writing 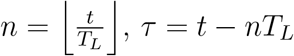, and *µ* = *A* + *C*, the method of steps [14, 25] gives

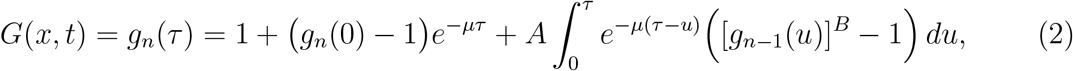

with 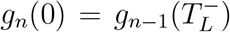. The full method-of-steps derivation of the analytical solution (2) is given in Appendix A. In practice, direct numerical integration of the delay differential equation (1) agrees with Eq. (2) to within numerical tolerance (maximum absolute error ∼ 3 × 10^−7^ across three independent solvers shown in Figure 4), so we use the method-of-steps recursion (2) in the remainder of the paper. In particular, the probability that a lineage has no free virions at time *τ* gives the *growth-only extinction* probability

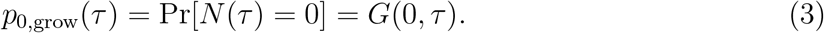

#### Basic reproduction number for free virion

A free virion either fails to infect before removal, with probability *C*/(*A* + *C*), or infects successfully, with probability *A*/(*A* + *C*), in which case it produces *B* offspring virions. The corresponding offspring generating function is 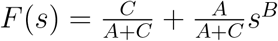. Its mean offspring number is

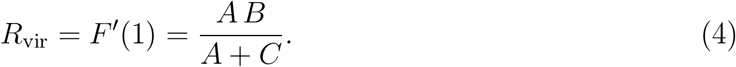

Thus *R*_vir_ is the mean offspring number of the free-virion branching process and serves as its basic reproduction number. Defining 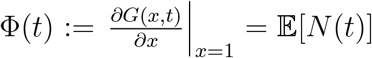 the expected number of free virions at time *t* in a lineage, yields

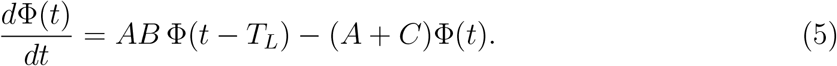

Seeking exponential solutions Φ(*t*) ∼ *e*^*λt*^, where *λ* denotes the early exponential growth rate, yields the characteristic equation

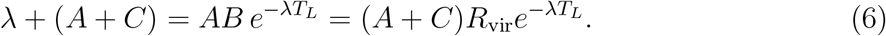

Setting *λ* = 0 in (6) gives the threshold condition *R*_vir_ = 1. Hence the process is supercritical if and only if *R*_vir_ > 1, in which case *λ* > 0.

#### Ultimate extinction probability from early exponential growth

Let *q* denote the probability of ultimate extinction of a lineage founded by a single free virion. By standard branching-process theory, *q* is the smallest root in [0, 1] of the fixed-point equation *s* = *F* (*s*), where 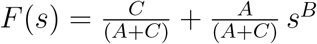. Equivalently, *q* satisfies

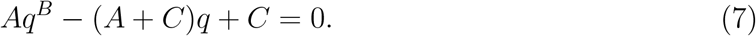

Recalling *F* ^′^(1) = *R*_vir_ in (4), we have that *q* = 1 if *R*_vir_ ≤ 1, then there exists a unique root *q* ∈ (0, 1), if *R*_vir_ > 1. Then the lineage survives with probability 1 − *q*. We emphasize that *q* is an ultimate extinction probability.

#### Extinction after growth and sampling

Suppose viral growth occurs for time *τ* followed by a sampling phase in which each virion survives independently with probability *D* (the bottleneck ratio). The sampling pgf is

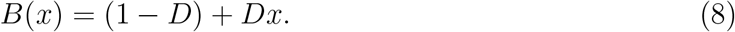

If *N*(*τ*) = *n*, the number of surviving virions is Binomial(*n, D*) with pgf ((1 − *D*) + *Dx*)^*n*^. Taking expectation over *N* (*τ*), the pgf of the surviving virion count after growth and sampling is

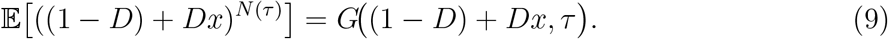

Evaluating at *x* = 0 gives the extinction probability after growth and sampling,

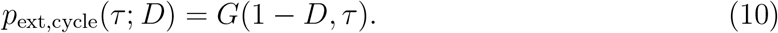

### 2.2 Parameterization of model (1) from the guinea-pig influenza data

With the stochastic framework in place, we now connect the model parameters to the guinea pig influenza data. From the viral titer data of [15], the parameters (*A, B, C*) are not jointly identifiable, since the characteristic equation (6) involves *A, B*, and *C* only through the combinations *AB* and *µ* = *A* + *C*, so distinct triples can produce the same early exponential slope *λ*. We therefore adopt a biologically constrained estimation strategy in which *T*_*L*_, *B*, and *C* are fixed to independent biological data (below), and *A* is then recovered by solving (6) for the unique value compatible with the observed *λ*,

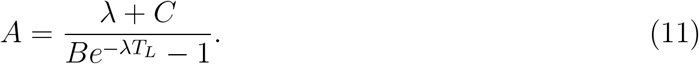

This yields a parameterization of (1) that is biologically interpretable and consistent with the limited information contained in early viral growth data.

#### Intracellular eclipse (latent) time *T*_*L*_

It denotes the interval between successful infection of a host cell and the first release of progeny virions into the free-virion compartment. Because guinea-pig nasal-wash samples were collected only once per day, the data do not resolve intracellular delays on the scale of hours [15]. We therefore fix *T*_*L*_ using published estimates of the influenza replication cycle from cell-culture and within-host studies. Influenza eclipse times are generally reported to be on the order of several hours, with a commonly cited range of approximately 5 − 12 hpi [5]. Specific estimates include a delay of about 6 h in human influenza kinetic fits [3] and mean eclipse phases of 6.6 h for wild type (WT) and 9.1 h for a mutant in MDCK cells [28]. We therefore take *T*_*L*_ = 6 h = 0.25 days as the nominal value and assess robustness over *T*_*L*_ ∈ [5, 12] h = [0.21, 0.5] days.

#### The burst size *B*

In this model, *B* denotes the total number of infectious progeny virions released by a productively infected cell over its lifespan. Because the free-virus state variable *N* (*t*) is interpreted as infectious virus in PFU-like units, *B* must likewise be interpreted as an *infectious burst size* (PFU/cell), rather than as the total number of physical particles produced per cell.

This interpretation is consistent with the experimental readout in the guinea pig study [15], where virus recovered by nasal lavage was quantified by plaque assay in MDCK cells and reported in PFU. However, the nasal-lavage measurements do not provide a direct in vivo estimate of per-cell burst size, so we fix *B* using strain-matched single-cycle measurements from cell culture.

For influenza A/Panama/2007/99 (H3N2), Jacobs *et al*. [18] reported that MDCK cells infected under single-cycle conditions produced a maximum of approximately 11.5 PFU per HA^+^ cell, with mean ± SD = 8.1–14.9 PFU/cell [18]. Moreover, Holmes *et al*. [15] found that the barcoded Pan/99 NA-BC virus used in the guinea pig transmission experiments showed no significant difference in MDCK growth kinetics relative to Pan/99 WT, supporting the use of the Pan/99 WT single-cycle estimate for this system. Accordingly, we adopt *B* = 10 PFU/cell as a nominal value with a range of [5, 20] PFU/cell.

#### The clearance rate *C*

The free-virus clearance rate *C* is related to the viral half-life *t*_1/2_ by 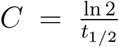, or equivalently, when *t*_1/2_ is measured in hours, 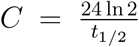 (day^−1^). Empirical influenza kinetic studies estimate early free-virus half-lives on the order of a few hours. Reported estimates include approximately 4 h in mice [24], 3.2 h in experimentally infected humans when an eclipse-phase model is used [3], and 2.3 h in a population model of human H1N1 infection [7]. These correspond to clearance rates of approximately *C* ≈ 4.2, 5.2, and 7.2 day^−1^. Accordingly, we consider *C* = 2.4–7.2 day^−1^, with a nominal value *C* = 4.8 day^−1^. This range corresponds to viral half-lives of approximately 7–2.3 hours and is broadly consistent with mammalian influenza kinetic studies as well as guinea pig infection experiments in which viral titers typically rise until about day 2 post infection.

##### 2.2.1 Estimation of the intrinsic early exponential growth rate

With the biological parameters fixed, the remaining quantity is the early growth rate *λ*. We estimate it from donor viral titers. In Eq. (5), *λ* is the dominant early exponential growth rate and reflects within-host replication kinetics once infection is established. Donor animals begin with a large founding population, so their aggregate titer trajectories are relatively insensitive to lineage-level stochastic loss. Barcode diversity also remains high in these animals, whereas recipient trajectories are strongly shaped by stochastic filtering after transmission.

During the early growth phase the titer *V* (*t*) satisfies *V* (*t*) ≈ *V*_0_*e*^*λt*^, so ln *V* (*t*) = ln *V*_0_ +*λt*. Since titers are reported as log_10_(PFU/mL), fitting log_10_ *V* (*t*) = *α* + *st* gives *λ* = *s* ln 10. Hence *λ* is directly estimable from the slope of the early log-linear rise in titer.

We used the log_10_ PFU/mL measurements of the 24 inoculated animals from the Holmes *et al*. S2 Data file [15], restricted to the earliest increasing interval, day 1 to day 2 post-infection, to minimize bias from later density dependence or immune control. The day-0 inoculation dose (5 × 10^4^ PFU) was excluded because it represents the administered inoculum rather than a within-host titer on the post-establishment growth trajectory and may include non-productive or rapidly cleared virions. For each animal *i, λ*_*i*_ = ln 10 [log_10_ *V*_*i*_(2) − log_10_ *V*_*i*_(1)], with *V*_*i*_(*t*) the titer on day *t*. Since the two observations are one day apart, *λ*_*i*_ has units of day^−1^. The resulting distribution is shown in Fig. 1 (Left). The median growth rate was 2.55 day^−1^ (Q1–Q3 1.64–3.36 day^−1^), corresponding to a doubling time *t*_double_ = ln 2/*λ* ≈ 0.27 days (≈ 6.5 h). Most estimates cluster within a biologically plausible range, but one outlier at 9.53 day^−1^ stands well apart from the remainder. As discussed in the next subsection, this value violates the stability condition required for the mechanistic inversion and was excluded from that step, leaving a maximum retained value of 6.28 day^−1^.

**Figure 1:**
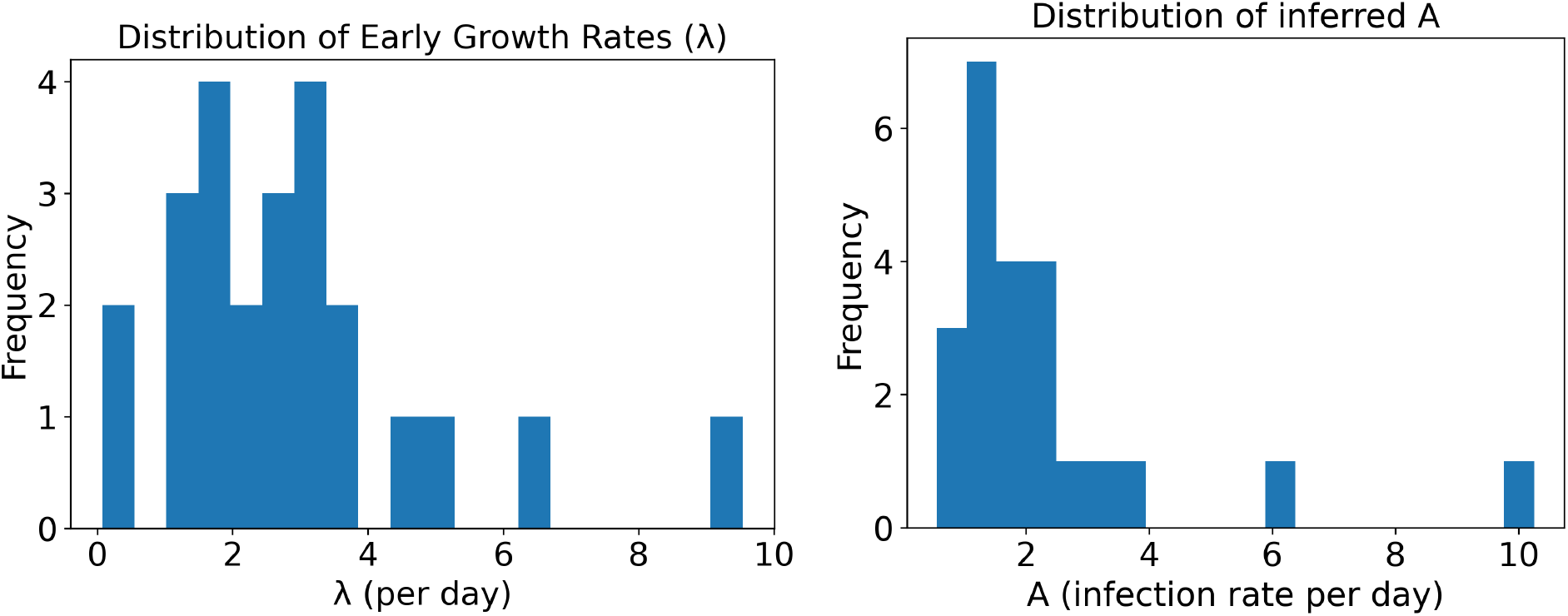
Left: Distribution of estimated early growth rates *λ* from all *n* = 24 inoculated animals. Right: Distribution of inferred infection rates *A* obtained by inversion of Eq. (11) from the *n* = 23 stability-filtered inoculated animals.

##### 2.2.2 Stability-constrained inversion to infection rate

Having estimated *λ*, we now invert the characteristic equation (6) to recover the infection-rate parameter *A* in (11). For biologically motivated parameters

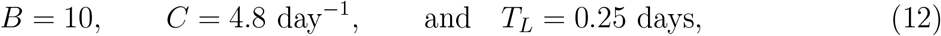

which apply to Figures 1–3 and Table 2 that follow, the inversion is well-defined only when 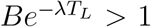, which defines the critical growth rate 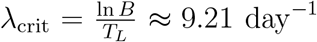. The maximal inferred value 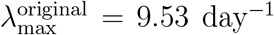 exceeds this stability threshold and therefore violates the dynamical feasibility condition. This single unstable estimate was excluded prior to mechanistic inversion, yielding a filtered set of *n* = 23 admissible growth rates with 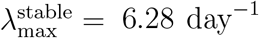. Per-animal estimates of the early growth rate *λ*_*i*_, the corresponding doubling time *t*_double,*i*_ = ln 2/*λ*_*i*_, and the inferred infection rate *A*_*i*_ are listed in Table 7 in Appendix B. The resulting distribution of infection rates is shown in Fig. 1 (Right). The inferred values were strictly positive, with median(*A*) = 1.6964 day^−1^ (Q1–Q3 1.1374–2.4014 day^−1^).

**Table 1:** Glossary of technical terms.

| Term | Definition |
| --- | --- |
| genetic diversity | The amount of genetic variation contained within a population. Can be described in terms of abundance and evenness. |
| transmission<br>bottleneck | The reduction in population size that occurs during transmission from a donor to recipient host. |
| adaptation | Evolution that results in an individual that is better fit in its current environment. |
| genetic drift | Loss of variants by chance, regardless of fitness. |
| genetic bottleneck | When a population experiences a dramatic reduction of genetic diversity, often caused by a dramatic reduction in population size, e.g. a small founding population from which a new infection grows. |
| virion | A single complete virus particle outside of a cell. |
| evolution | Descent with inherited modification. |
| mutation | A small change in the genetic code, often arising from a random copying error during replication. |
| genotype | The complete set of genetic material. |

Table 1 continued from previous page.
| Term | Definition |
| --- | --- |
| antigenic variation | A defense strategy employed by pathogens, including viruses, where they vary their surface structures to avoid detection by the host immune system. Host immune systems can recognize surface structures of pathogens, e.g. through antibodies, so pathogens that sufficiently alter their surface structures can then evade detection by host immune cells. |
| fitness | The reproductive success of a variant. Variants with higher fitness leave more progeny in a given environment. |
| recipient-side filter | Recipient airway processes that remove free virions beyond what stochastic growth predicts. Examples include superinfection exclusion, target-cell competition, and mucociliary transport. |
| stochastic dynamics | Chance fluctuations in viral lineage numbers during early replication that can wipe out individual lineages. |
| positive selection | Selection that favors trait values at one extreme, shifting the distribution mean toward that extreme. |
| purifying selection | Selection that favors intermediate trait values over extremes, reducing the variance of the trait distribution. |

**Table 2:**
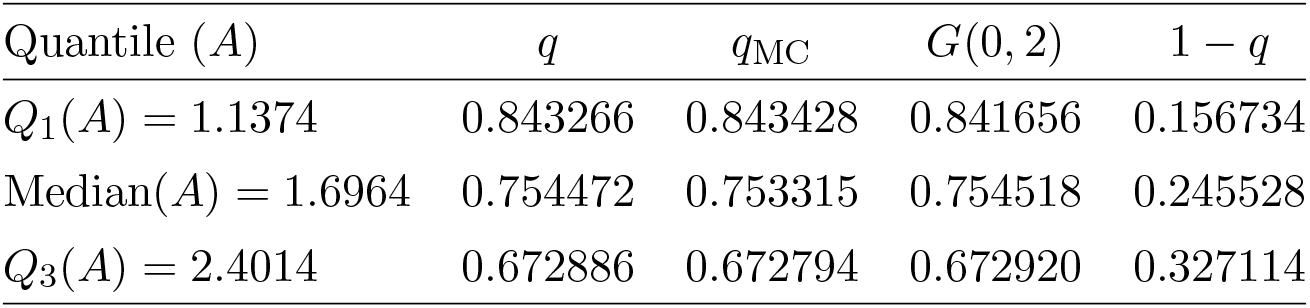
Asymptotic versus finite-time growth-only extinction probabilities.

#### Nonlinearity of the *λ* ↦ *A* map

The map *λ* ↦ *A*(*λ*) (Fig. 2 (Left)) is monotone increasing and strongly nonlinear. Differentiating (11) by the quotient rule gives

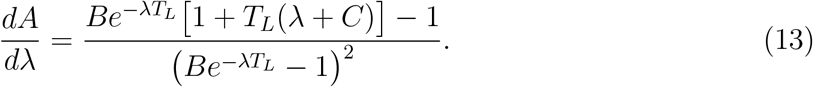

**Figure 2:**
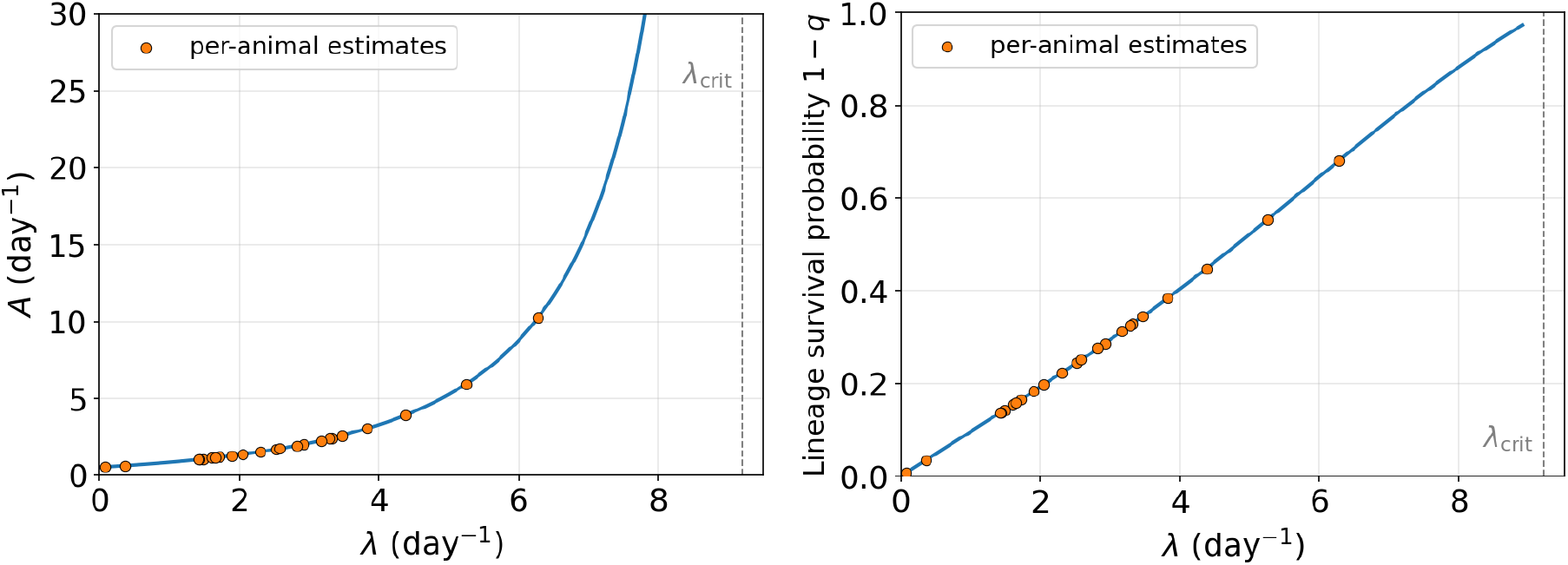
Left: Map from empirical growth rate *λ* to mechanistic infection rate *A* (Eq. (11)). Right: Composite map from empirical growth rate *λ* to ultimate lineage survival probability 1 − *q*. Orange points are the 23 per-animal estimates.

As *λ* → *λ*_crit_ the denominator 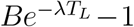 in (11) vanishes and *dA*/*dλ* diverges. We emphasize that *λ*_crit_ is an *upper feasibility limit* imposed by the finite latent period and burst size, not the branching-process supercriticality threshold, which is *R*_vir_ = 1, equivalently *λ* = 0. The largest admissible estimate is 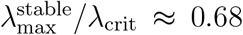, so the divergence itself is not reached. The monotone nonlinearity of the map therefore propagates the wide range of empirical growth rates (*λ* ≈ 0.07–6.28 day^−1^) into a correspondingly wide range of infection rates (*A* ≈ 0.55–10.25 day^−1^).

Because extinction probabilities in the branching framework depend on *A*, not on *λ*, the nonlinear inversion propagates growth-rate heterogeneity into heterogeneous lineage survival probabilities. Figure 2 (Right) shows the composite map *λ* ↦ 1 − *q*, obtained by composing the stability-constrained inversion *λ* ↦ *A* (Eq. (11)) with the fixed-point equation for *q* (Eq. (7)), with the 23 per-animal estimates overlaid. This composite map admits a closed form. Since *q* = 1 is always a root of (7) (its coefficients sum to zero), we factor it out using 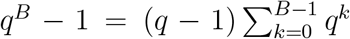, so that the nontrivial survival root *q* ∈ (0, 1) satisfies 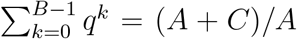. Substituting the inversion (11) for *A* eliminates the infection rate and yields

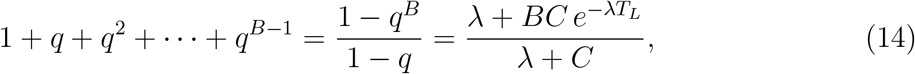

which defines *q* = *q*(*λ*) implicitly, with lineage survival probability 1 − *q*. For *B* = 10 the surviving root solves a degree-nine polynomial with no radical closed form and is evaluated numerically. Notably, the composite *λ* ↦ 1 − *q* is approximately linear over the observed range (Fig. 2, Right). Across this range, lineage survival spans 1−*q* ≈ 0.007–0.68, a ∼95-fold range.

#### Propagation of parameter uncertainty to extinction dynamics

To quantify how un-certainty in the inferred infection rate propagates into lineage survival, we evaluated the finite-time growth-only extinction probability *p*_0,grow_(*t*) (Eq. (3)) for the lower-quartile, median, and upper-quartile values of *A* obtained from the stability-filtered *λ* distribution. As shown in Fig. 3, all three trajectories rise rapidly during the first eclipse period. Before the first burst (*t* < *T*_*L*_) no offspring have yet been produced, so *p*_0,grow_(*t*) = 1 − *e*^−(*A*+*C*)*t*^ increases monotonically to a peak 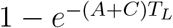 at *t* = *T*_*L*_, equal to 0.77, 0.80, and 0.83 for *Q*_1_, the median, and *Q*_3_ of *A*, respectively. Thereafter the curve relaxes toward *q*. A transient overshoot therefore appears whenever this peak exceeds *q*. It is absent for *Q*_1_ (peak 0.77 < *q* = 0.84, monotone rise) but present for the median (0.80 > *q* = 0.75) and *Q*_3_ (0.83 > *q* = 0.67). This highlights the distinction between the finite-time quantity *p*_0,grow_(*t*), the probability that no free mutant virions are present at time *t*, and the ultimate extinction probability *q*, the asymptotic probability of permanent extinction. Table 2 reports, for each quartile, the theoretical *q* from Eq. (7), the Monte Carlo estimate *q*_MC_ computed from 10^6^ independent simulations with a fixed random seed, and the endpoint *G*(0, 2) of the Fig. 3 trajectories, which closely match *q* and confirm that the finite-time curves have essentially converged by *t* = 2 days. Increasing *A* lowers *q* and raises the lineage-survival probability 1 − *q*. The spread between quartiles therefore measures how uncertainty in *A* propagates into uncertainty in early lineage survival.

**Figure 3:**
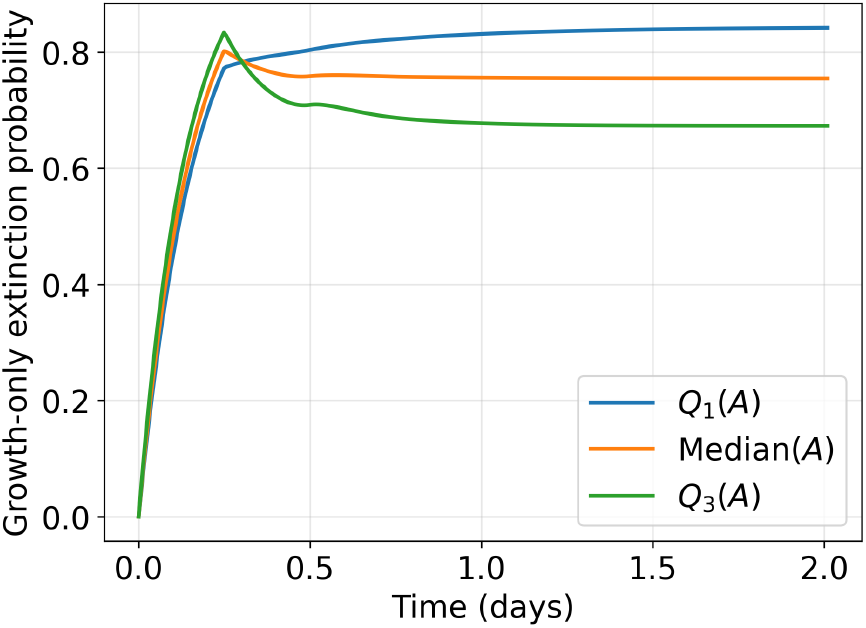
Finite-time growth-only extinction probability *p*_0,grow_(*t*) (Eq. (3)) for the infection-rate quartiles *Q*_1_ = 1.1374, Median = 1.6964, and *Q*_3_ = 2.4014 day^−1^. Each trajectory converges to the corresponding ultimate extinction probability *q* (Eq. (7)).

**Figure 4:**
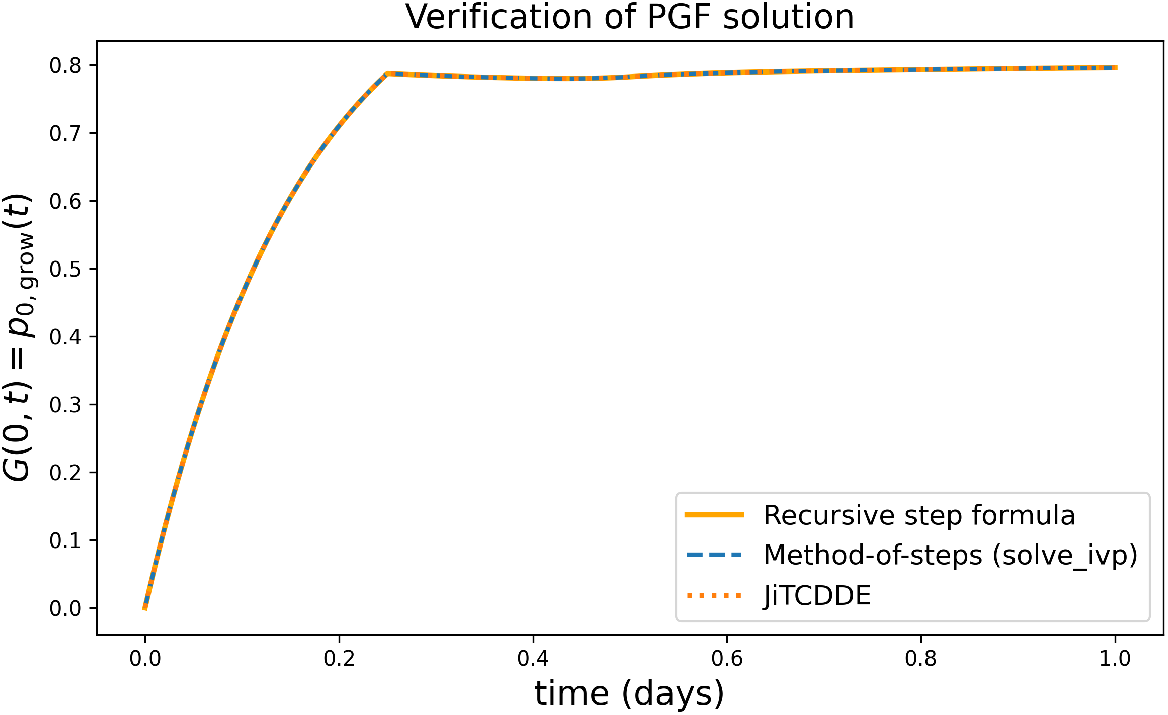
Numerical verification of the finite-time PGF solution. The recursive step formula for *G*(0, *t*) is compared with two independent numerical integrations of (1). The three curves are visually indistinguishable.

## 3 Empirical lineage contraction in recipient animals

Before modeling recipient-side lineage loss, we note that the diversity collapse observed in recipient animals cannot be attributed to stochastic sampling at inoculation. In the guinea pig experiments of [15], each donor was inoculated intranasally with *N*_tot_ = 5 × 10^4^ plaque-forming units (PFU) of the Pan/99 NA-BC virus, distributed across *N*_bar_ effective barcode lineages. Quantifying the effective lineage count by the exponential of the reported Shannon diversity (*H* = 7.<u>90 ±</u> 0.02) gives *N*_bar_ ≈ *e*^*H*^ ≈ 2600–2800, so that each lineage is represented on average by 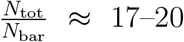 infectious virions. With this level of redundancy, undersampling at the inoculation step alone cannot produce the sharp contractions observed in recipients. The diversity loss must therefore originate *after* transmission, during early viral expansion in the host. We model this process in the remainder of the section using the branching framework of Section 2 with the parameter estimates of Section 2.2, augmented by an explicit recipient-side filter that captures losses in the establishment stage not accounted for by intrinsic stochasticity alone.

### 3.1 Experimental design and data source

The barcode data of Holmes et al. [15] span three independent transmission experiments (E1, E2, E3), each consisting of inoculated donor animals paired with naïve contact (“exposed”) recipients. Donors were inoculated intranasally with 5 × 10^4^ PFU of the Pan/99 NA-BC barcoded virus on day 0. Contact recipients were co-housed beginning on day 1. Nasal-wash samples were collected daily from each animal and subjected to amplicon sequencing of the 12-site barcode locus. Holmes et al. report Shannon diversity, observed richness, and evenness for each sample, together with 95% confidence intervals obtained by subsampling the barcode reads at multiple depths.

In total, the three experiments yield 24 recipient animals with at least two plaque-positive days of sequencing data (Table 8 in Appendix B). For each recipient, we define *t*_1_ = first plaque-positive day (days post inoculation of the donor) and *t*_2_ = second plaque-positive day. In all 24 cases the interval *t*_2_ − *t*_1_ equals one calendar day.

### 3.2 Estimating the effective number of lineages and empirical contraction ratio

Raw barcode counts overestimate the number of biologically meaningful lineages because many barcodes appear at very low frequency and may represent sequencing artefacts or transient detection. We therefore quantify the effective number of lineages using the *Hill number of order 1*, defined as

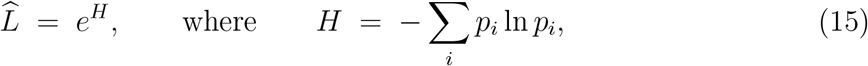

*p*_*i*_ is the relative frequency of barcode *i* in the sample, and *H* is the *Shannon entropy* (also called *Shannon diversity*). This measure gives the number of equally abundant lineages that would produce the same Shannon entropy, and is less sensitive to rare barcodes than observed richness. For each recipient we compute 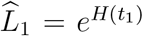 and 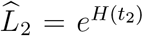, and define the *empirical contraction ratio*

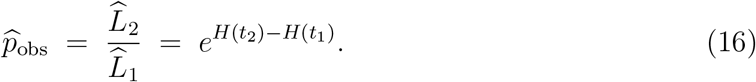

Equation (16) shows that the contraction ratio 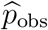 (left) equals both the ratio of Hill numbers 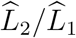 (middle) and the exponential of the Shannon-entropy difference (right). The contraction can therefore be read directly from the change in Shannon diversity *H*(*t*_2_) − *H*(*t*_1_) without exponentiating each value separately.

#### Uncertainty quantification

Holmes et al. report 95% confidence intervals (CI) for Shannon diversity, obtained by subsampling barcode reads at multiple depths. The lower and upper bounds are provided as columns `Shannon_low` and `Shannon_high` in their supplementary data. We denote these *H*^low^ and *H*^high^ and propagate them into conservative bounds on 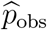 by taking the worst-case combination as 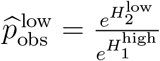 and 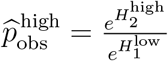. This yields the widest plausible range for 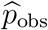. The resulting 95% CIs are reported in the last column of Table 8.

### 3.3 Empirical contraction patterns

The per-animal values of 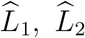, and 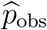 are listed in Table 8 (Appendix B). The key findings, summarized in Table 3, are as follows.

**Table 3:** Summary of the empirical contraction ratio 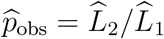 and the effective number of lineages 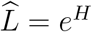 (Shannon-based Hill number) by experiment. *n* is the number of recipients.

| Experiment | $n$ | $\hat{L}_1$ | $\hat{L}_2$ | $\hat{p}_{\text{obs}}$ |
| --- | --- | --- | --- | --- |
|  |  | median [Q1, Q3] | median [Q1, Q3] | median [Q1, Q3] |
| E1 | 8 | 792.1 [366, 822] | 6.3 [4.3, 55] | 0.011 [0.005, 0.224] |
| E2 | 8 | 99.8 [90, 103] | 76.7 [42, 98] | 0.860 [0.391, 1.094] |
| E3 | 8 | 42.5 [39, 74] | 8.8 [4.7, 33] | 0.162 [0.104, 0.472] |
| All | 24 | 100.6 [47, 265] | 33.3 [4.8, 75] | 0.219 [0.040, 1.071] |

#### Experiment 1: severe contraction

E1 recipients begin with very high effective diversity 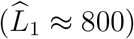, consistent with a large and diverse inoculum successfully seeding many lineages. By the second positive day, diversity collapses to single digits 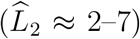 for five of eight animals, yielding 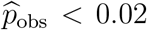. The exception is GP14, for which 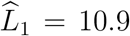 on day 2 but 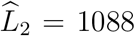 on day 3, giving 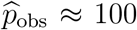. This animal was likely barely positive on its first sampling day, with only a few dominant lineages detectable. By the following day the full diverse inoculum from the donor had become established. GP14 therefore represents an expansion rather than a contraction and is excluded from the contraction analysis.

#### Experiment 2: bimodal behavior

E2 recipients start at lower diversity 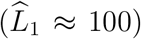. Four animals (GP25–28) show moderate to strong contraction 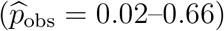, while the other four (GP29–32) maintain or slightly increase diversity 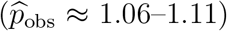. The latter group is consistent with growth-only dynamics in which all initially detected lineages persist.

#### Experiment 3: heterogeneous contraction

E3 exhibits intermediate and heterogeneous behavior. Six of eight recipients show clear contraction 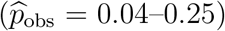, while two (GP40, GP44) show mild expansion 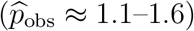.

#### Overall pattern

Across all 24 recipients, the median contraction ratio is 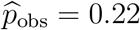 (Q1– Q3 0.04–1.07), indicating that the typical recipient retains roughly one-fifth of the effective lineage diversity observed on the first positive day. The wide spread reflects the heterogeneity across experiments and animals. The corresponding contraction strength varies over more than two orders of magnitude, from 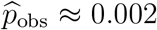 in the most severely contracting recipient to ≈ 1.6 in the least (excluding one expansion outlier; Table 8). These observations raise a natural question. For animals with 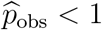, can the observed loss be explained by growth-only extinction, or does it require an additional recipient-side filter?

### 3.4 Growth-only baseline and classification

To determine whether the observed contraction can be explained by stochastic growth alone, we compare 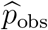 to the growth-only baseline. Under growth-only dynamics, a single founder virion is represented in the free-virion pool at time *t* with probability 1 − *G*(0, *t*), where *G*(0, *t*) is the growth-only extinction probability from Eq. (3).

Table 4 summarizes the growth-only survival probabilities at the three representative infection rates from Section 2.2.

**Table 4:**
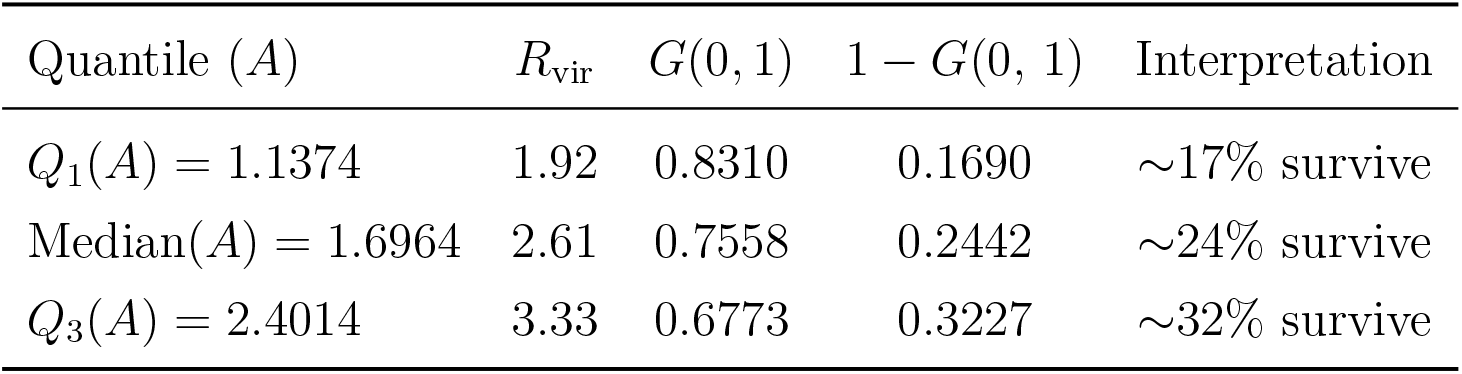
Growth-only baseline for three *A* quartiles at *t* = 1 day. Parameters from Eq. (12).

| Quantile ($A$) | $R_{\text{vir}}$ | $G(0, 1)$ | $1 - G(0, 1)$ | Interpretation |
| --- | --- | --- | --- | --- |
| $Q_1(A) = 1.1374$ | 1.92 | 0.8310 | 0.1690 | $\sim 17\%$ survive |
| Median($A$) = 1.6964 | 2.61 | 0.7558 | 0.2442 | $\sim 24\%$ survive |
| $Q_3(A) = 2.4014$ | 3.33 | 0.6773 | 0.3227 | $\sim 32\%$ survive |

Even under supercritical branching with *R*_vir_ > 1, only 17 to 32% of single-virion founders are represented in the free-virion pool after one day, as shown in Table 4. This reflects two factors. The high clearance rate *C* = 4.8 day^−1^ gives a viral half-life of approximately 3.5 hours. And a single founder faces stochastic extinction before producing offspring at the first burst *t* = *T*_*L*_ = 6 hours.

The empirical contraction ratio 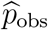 can therefore be compared against this baseline. If 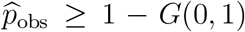, growth-only dynamics suffice. If 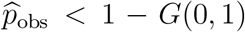, the observed diversity collapse exceeds what stochastic extinction alone can explain, and an additional recipient-side filter is required.

Applying this test at the median infection rate *A* = 1.6964 day^−1^, with baseline 1 − *G*(0, 1) = 0.244, the 24 recipients fall into three groups. Twelve animals have 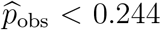 and are classified as *filter needed*, with observed contraction exceeding the growth-only prediction. Five animals satisfy 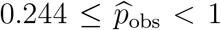 and are classified as *growth-only sufficient*, meaning that intrinsic branching extinction alone accounts for the observed loss. Six animals show 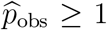 and are classified as *no contraction*, with diversity maintained or increasing from *t*_1_ to *t*_2_. The remaining animal GP14 is an expansion outlier and is excluded from the analysis. This classification uses the median *A* as the canonical infection rate. At higher *Q*_3_(*A*), the baseline 1 − *G*(0, 1) shifts upward, and one additional animal, GP48 in E3, would be reclassified as *filter needed*.

## 4 Establishment filtering and the emergent recipient bottleneck

### 4.1 Filtered branching-process framework

During establishment in the recipient, free virions may be removed by recipient-side processes beyond what the growth-only dynamics already account for, including competition for susceptible target cells [3, 29], superinfection exclusion that renders recently infected cells refractory to coinfection [30], spatial exclusion from target-cell niches [18], and mucociliary transport. These processes act continuously as the population grows, but we approximate their combined effect by a single filtering step applied to the population after growth. This approximation suits two of the four mechanisms especially well. Competition for target cells and superinfection exclusion are strongest when the free-virion and infected-cell populations are large, that is, late in the interval once the population has expanded, so applying their effect to the grown population captures most of the loss. Mucociliary clearance acts more evenly over time, but across the short Δ*t* = 1 day window its net effect is still well summarized by a single survival fraction.

We therefore reuse the growth-and-sampling construction of Eqs. (8)–(9), applied to the recipient with an effective per-virion survival probability 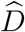. The model does not claim that removal physically occurs at a single instant. Rather, the net cumulative effect of these continuous processes on a free virion’s chance of founding a persisting lineage is summarized by one survival probability 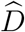, applied to the grown free-virion pool at the readout time. “After growth” is the observation point, not a statement about when removal acts.

For a single founder lineage with *N*_*t*_ free-virion descendants at time *t*, specializing the growth-and-sampling extinction probability Eq. (10) to *τ* = *t* and 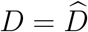, and writing 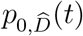 for the result, gives the probability that *no* descendant survives both growth and filtering,

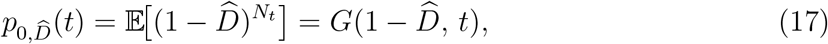

so the lineage is represented at time *t* with probability 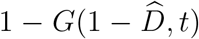. Here 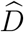 is a per-descendant pass probability, not a lineage-level establishment probability. The lineage-level survival 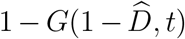 is a nonlinear function of 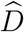, mediated by the full distribution of *N*_*t*_. This factored form is *exact* when each free virion is removed independently of the others. Independence is what lets us treat the survivors as a 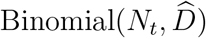 draw, which gives the sampling pgf of Eq. (9) and, at *x* = 0, reduces to Eq. (10). In practice the removal need not be fully independent. Virions descended from the same founder tend to lie close together, so a spatially localized clearance event such as a focal immune response can remove many of them at once. When removal is correlated in this way, 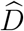 no longer represents a literal per-virion survival probability and is better understood as an effective parameter.

A single effective filter is also the most the present data can support. Each recipient contributes only two diversity snapshots one day apart, which constrain a single net survival parameter but cannot resolve the temporal profile of removal. A more finely time-resolved model would be unidentifiable, so 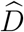 is the simplest choice the data can support.

Two boundary cases clarify the interpretation. When 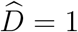 no additional filter is present and the model reduces to growth-only extinction. When 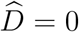 every descendant is removed and extinction is certain.

### 4.2 Linking the contraction ratio to the filter

To estimate the filter from the data, we link the contraction ratio 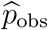 to 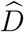 over the one-day observation interval Δ*t* = *t*_2_ − *t*_1_. We reset the branching process at *t*_1_ and treat each of the 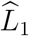 lineages as a single effective founder that independently undergoes branching and filtering over Δ*t*. Fixing Eq. (10) to *τ* = Δ*t* and 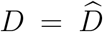, each founder survives with probability 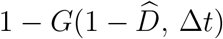, giving

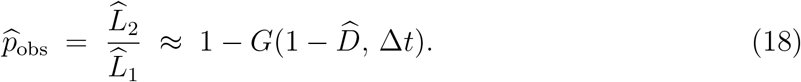

Resetting at *t*_1_ is justified because by then the growth-only extinction probability *p*_0,grow_(*t*) = *G*(0, *t*) from Eq. (3) has approximately converged to the ultimate extinction probability *q* from Eq. (7). All 24 recipients have Δ*t* = 1 day and the observation times *t*_1_, *t*_2_ are well beyond the eclipse period *T*_*L*_ = 0.25 days. Recipients are co-housed from donor day 1, so the branching process in each recipient has been running for at most *t*_1_ − 1 days by the first observation, approximately one day for the majority of animals with *t*_1_ = 2. The *G*(0, 1) values are tabulated in Table 4, and the corresponding *q* values in Table 2. The relative differences in 1 − *q* are 0.5%, 8%, and 1.3% for the median, *Q*_1_, and *Q*_3_ values of *A*, so the approximation is best near the median *A* and looser at the extremes.

The branching-process model assumes that each barcode lineage in the recipient is founded by a single free virion. If a lineage is instead seeded by *m* > 1 virions, its true survival probability is 1 − *q*^*m*^ rather than 1 − *q*, and the model overestimates growth-only extinction. The bias would push 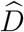 upward, making the estimated filter appear more permissive than it is. Two features limit this bias in our setting. First, contact-transmitted recipients receive a small per-lineage founding dose. Second, the Hill-number ratio 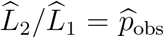 from Eq. (16) depends only on the relative frequencies of lineages, and a common multi-founder scaling preserves those frequencies. The multi-founder bias affects absolute survival but not the shape captured by 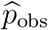, so it does not propagate into 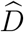. In systems with a larger per-lineage inoculum, the multi-founder generalisation 1 − *q*^*m*^ should be used in place of 1 − *q*.

### 4.3 Estimated filter strength and sensitivity

#### 4.3.1 Per-animal filter estimates

For the 12 recipients requiring an additional filter, we rearrange Eq. (18) as

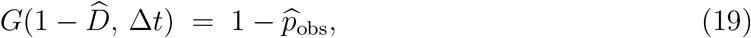

and solve for 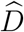 by bisection on *D* ∈ (0, 1], using the method-of-steps recursion for *G*(*x, t*) from Appendix A. Uncertainty in 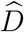 from the assumed infection rate is captured by the per-quartile estimates on *A*. Within each quartile of *A*, 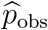 is held at its per-animal point estimate. As a result, sampling uncertainty in 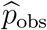 is not propagated. Table 5 summarizes the resulting 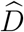 values: at each row’s *A*, we compute 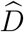 per animal and report the across-animal median with the Q1–Q3 bracket, with *n* the number of filter-needed animals with 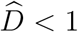 at that *A*. The full per-animal estimates are given in Table 9 in Appendix B. The estimated filter parameters are very small. At the median infection rate, the median 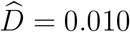 (Q1–Q3 0.002–0.033), meaning that roughly 1 in 100 free virions passes the recipient-side filter.

**Table 5:** Summary of 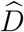 for filter-needed recipients.

| Quantile ($A$) | $n$ | median($\hat{D}$) | $Q_1(\hat{D})$–$Q_3(\hat{D})$ |
| --- | --- | --- | --- |
| $Q_1(A) = 1.1374$ | 11 | 0.0150 | [0.0038, 0.0556] |
| Median($A$) = 1.6964 | 12 | 0.0099 | [0.0018, 0.0328] |
| $Q_3(A) = 2.4014$ | 12 | 0.0047 | [0.0009, 0.0145] |

Table 5 pools all filter-needed animals at each *A*, so it captures sensitivity to the assumed infection rate but averages over experimental context. Because the three experiments are independent replicates with different donor and recipient cohorts, we report 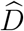 per experiment in Table 6 to expose between-experiment variation that the pooled summary hides. We fix *A* at its median value (1.6964 day^−1^), compute 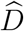 per animal, and report the within-experiment median with the Q1–Q3 bracket. Only filter-needed animals are included, and *n* is the number of such animals per experiment. E1 recipients experience the tightest filtering (median 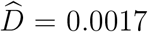, approximately 2 in 1000), while E3 recipients show a more moderate filter (median 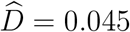, approximately 5 in 100).

**Table 6:** Per-experiment summary of 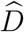 at the median infection rate *A*.

| Experiment | $n$ | median($\hat{D}$) | $Q_1(\hat{D})$–$Q_3(\hat{D})$ |
| --- | --- | --- | --- |
| E1 | 6 | 0.0017 | [0.0010, 0.0035] |
| E2 | 1 | 0.0065 | — |
| E3 | 5 | 0.0454 | [0.0286, 0.0644] |

**Table 7:** Animal-specific estimates of the early exponential growth rate *λ*, the associated doubling time *t*_double_, and the corresponding inferred infection rate *A* (Eq. (11), with *B, C*, and *T*_*L*_ as in the main text). The unstable case for animal 24 was excluded from inference of *A*. Only inoculated animals are included; IDs follow the original study’s numbering.

| Animal ID | $\lambda$ (day<sup>-1</sup>) | Doubling time (h) | $A$ (day<sup>-1</sup>) |
| --- | --- | --- | --- |
| 1 | 2.0437 | 8.14 | 1.3689 |
| 3 | 2.5257 | 6.59 | 1.6964 |
| 5 | 3.3242 | 5.00 | 2.4209 |
| 7 | 3.2879 | 5.06 | 2.3818 |
| 9 | 1.8971 | 8.77 | 1.2822 |
| 11 | 1.4827 | 11.22 | 1.0644 |
| 13 | 2.9223 | 5.69 | 2.0235 |
| 15 | 1.7207 | 9.67 | 1.1847 |
| 17 | 1.6094 | 10.34 | 1.1270 |
| 18 | 4.3820 | 3.80 | 3.9177 |
| 19 | 3.4657 | 4.80 | 2.5794 |
| 20 | 6.2791 | 2.65 | 10.2500 |
| 21 | 3.8250 | 4.35 | 3.0334 |
| 22 | 5.2495 | 3.17 | 5.9402 |
| 23 | 1.4318 | 11.62 | 1.0402 |
| 24 | 9.5337 | 1.74 | — |
| 33 | 0.3604 | 46.15 | 0.6341 |
| 35 | 0.0741 | 224.48 | 0.5528 |
| 37 | 3.1676 | 5.25 | 2.2572 |
| 39 | 2.5802 | 6.45 | 1.7380 |
| 41 | 2.8239 | 5.89 | 1.9368 |
| 43 | 1.6503 | 10.08 | 1.1478 |
| 45 | 2.3026 | 7.22 | 1.5362 |
| 47 | 1.4214 | 11.70 | 1.0353 |

**Table 8:** Per-animal recipient lineage counts and contraction ratios. *t*_1_, *t*_2_ are the first and second plaque-positive day (dpi of donor). 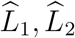 are Shannon-based effective lineage counts (*e*^*H*^) computed from Holmes’ Shannon diversity, with the values in parentheses giving the 95% CI endpoints 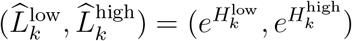, propagated from Holmes’ subsampling-based Shannon bounds. 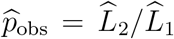 is the Hill-based contraction ratio. The 95% CI column gives its worst-case propagated bounds 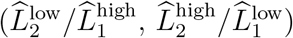. The last column *R*_2_/*R*_1_ is the raw-richness contraction ratio computed from Holmes’ observed-richness data, included as a sensitivity comparator. GP14 (marked ^∗^) is an expansion outlier excluded from the contraction analysis.

| Exp | GP | $t_1$ | $t_2$ | $\hat{L}_1$ ($\hat{L}_1^{\text{low}}, \hat{L}_1^{\text{high}}$) | $\hat{L}_2$ ($\hat{L}_2^{\text{low}}, \hat{L}_2^{\text{high}}$) | $\hat{p}_{\text{obs}}$ | $\hat{p}_{\text{obs}}$ 95% CI | $R_2/R_1$ |
| --- | --- | --- | --- | --- | --- | --- | --- | --- |
| E1 | 2 | 2 | 3 | 774.6 (762.2, 787.3) | 2.5 (2.4, 2.6) | 0.0033 | (0.0031, 0.0034) | 0.0626 |
| E1 | 4 | 2 | 3 | 809.6 (797.1, 822.4) | 4.9 (4.7, 5.1) | 0.0061 | (0.0058, 0.0064) | 0.0765 |
| E1 | 6 | 2 | 3 | 441.9 (430.0, 454.0) | 6.8 (6.6, 7.1) | 0.0155 | (0.0145, 0.0165) | 0.0891 |
| E1 | 8 | 2 | 3 | 814.8 (801.7, 828.2) | 5.8 (5.7, 6.0) | 0.0072 | (0.0069, 0.0075) | 0.1109 |
| E1 | 10 | 4 | 5 | 843.4 (830.5, 856.5) | 38.0 (37.4, 38.6) | 0.0451 | (0.0437, 0.0465) | 0.0526 |
| E1 | 12 | 3 | 4 | 1035.3 (1022.3, 1048.4) | 1.9 (1.9, 1.9) | 0.0018 | (0.0018, 0.0019) | 0.0225 |
| E1 | 14* | 2 | 3 | 10.9 (10.8, 11.0) | 1087.7 (1071.8, 1103.8) | 99.70 | (97.52, 101.94) | 24.06 |
| E1 | 16 | 4 | 5 | 139.9 (135.8, 144.2) | 106.5 (104.4, 108.6) | 0.7609 | (0.7241, 0.7997) | 0.8177 |
| E2 | 25 | 2 | 3 | 113.0 (110.2, 115.8) | 42.3 (41.4, 43.3) | 0.3746 | (0.3575, 0.3926) | 0.2951 |
| E2 | 26 | 2 | 3 | 99.2 (97.1, 101.3) | 2.4 (2.3, 2.5) | 0.0239 | (0.0223, 0.0256) | 0.2532 |
| E2 | 27 | 2 | 3 | 91.2 (87.9, 94.6) | 59.8 (57.5, 62.2) | 0.6559 | (0.6077, 0.7078) | 0.7398 |
| E2 | 28 | 2 | 3 | 100.3 (98.3, 102.4) | 39.8 (38.7, 40.9) | 0.3966 | (0.3785, 0.4156) | 0.2786 |
| E2 | 29 | 2 | 3 | 86.9 (82.9, 91.1) | 95.0 (93.6, 96.4) | 1.0929 | (1.0275, 1.1624) | 1.0586 |
| E2 | 30 | 2 | 3 | 100.8 (97.8, 103.9) | 107.2 (105.5, 109.0) | 1.0642 | (1.0153, 1.1154) | 1.0010 |
| E2 | 31 | 2 | 3 | 107.7 (105.5, 109.9) | 119.5 (117.2, 121.8) | 1.1094 | (1.0659, 1.1546) | 1.0367 |
| E2 | 32 | 2 | 3 | 85.4 (82.5, 88.3) | 93.7 (90.5, 97.0) | 1.0975 | (1.0247, 1.1753) | 1.0067 |
| E3 | 34 | 2 | 3 | 48.4 (47.6, 49.1) | 4.0 (3.9, 4.1) | 0.0823 | (0.0790, 0.0858) | 0.8489 |
| E3 | 36 | 2 | 3 | 33.8 (33.3, 34.3) | 4.6 (4.5, 4.6) | 0.1353 | (0.1316, 0.1391) | 1.7546 |
| E3 | 38 | 2 | 3 | 42.0 (41.6, 42.3) | 4.7 (4.5, 4.9) | 0.1115 | (0.1057, 0.1177) | 1.4291 |
| E3 | 40 | 2 | 3 | 42.9 (42.0, 43.9) | 69.3 (67.3, 71.3) | 1.6135 | (1.5354, 1.6956) | 1.6209 |
| E3 | 42 | 2 | 3 | 206.0 (199.7, 212.5) | 9.3 (9.0, 9.6) | 0.0449 | (0.0421, 0.0479) | 0.0766 |
| E3 | 44 | 2 | 3 | 40.1 (39.8, 40.4) | 45.5 (44.8, 46.2) | 1.1356 | (1.1107, 1.1612) | 1.4419 |
| E3 | 46 | 2 | 3 | 151.9 (146.7, 157.3) | 28.5 (28.3, 28.7) | 0.1878 | (0.1799, 0.1960) | 0.0973 |
| E3 | 48 | 2 | 3 | 33.1 (32.9, 33.3) | 8.3 (8.1, 8.5) | 0.2510 | (0.2432, 0.2590) | 1.2762 |

**Table 9:**
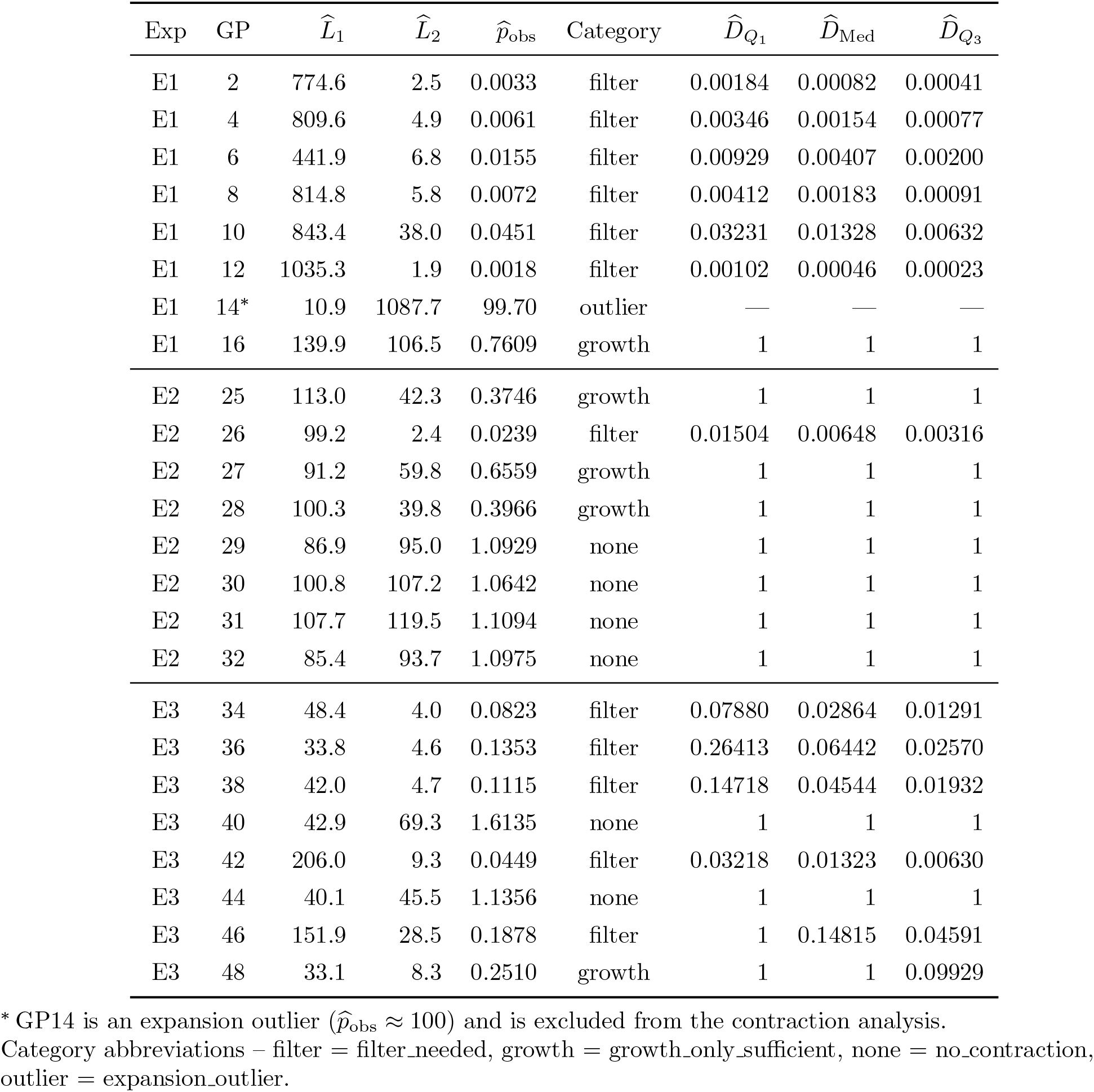
Per-animal 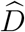 estimates under the effective-interval formulation. Baseline is 1 − *G*(0, 1) = 0.244 at median *A* (Table 4). 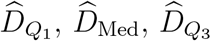 correspond to *A* = 1.1374, 1.6964, 2.4014 day^−1^, respectively. A dash (—) indicates the estimate is not applicable. 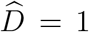 indicates no additional filter is needed at that *A*.

| Exp | GP | $\hat{L}_1$ | $\hat{L}_2$ | $\hat{p}_{\text{obs}}$ | Category | $\hat{D}_{Q_1}$ | $\hat{D}_{\text{Med}}$ | $\hat{D}_{Q_3}$ |
| --- | --- | --- | --- | --- | --- | --- | --- | --- |
| E1 | 2 | 774.6 | 2.5 | 0.0033 | filter | 0.00184 | 0.00082 | 0.00041 |
| E1 | 4 | 809.6 | 4.9 | 0.0061 | filter | 0.00346 | 0.00154 | 0.00077 |
| E1 | 6 | 441.9 | 6.8 | 0.0155 | filter | 0.00929 | 0.00407 | 0.00200 |
| E1 | 8 | 814.8 | 5.8 | 0.0072 | filter | 0.00412 | 0.00183 | 0.00091 |
| E1 | 10 | 843.4 | 38.0 | 0.0451 | filter | 0.03231 | 0.01328 | 0.00632 |
| E1 | 12 | 1035.3 | 1.9 | 0.0018 | filter | 0.00102 | 0.00046 | 0.00023 |
| E1 | 14* | 10.9 | 1087.7 | 99.70 | outlier | — | — | — |
| E1 | 16 | 139.9 | 106.5 | 0.7609 | growth | 1 | 1 | 1 |
| E2 | 25 | 113.0 | 42.3 | 0.3746 | growth | 1 | 1 | 1 |
| E2 | 26 | 99.2 | 2.4 | 0.0239 | filter | 0.01504 | 0.00648 | 0.00316 |
| E2 | 27 | 91.2 | 59.8 | 0.6559 | growth | 1 | 1 | 1 |
| E2 | 28 | 100.3 | 39.8 | 0.3966 | growth | 1 | 1 | 1 |
| E2 | 29 | 86.9 | 95.0 | 1.0929 | none | 1 | 1 | 1 |
| E2 | 30 | 100.8 | 107.2 | 1.0642 | none | 1 | 1 | 1 |
| E2 | 31 | 107.7 | 119.5 | 1.1094 | none | 1 | 1 | 1 |
| E2 | 32 | 85.4 | 93.7 | 1.0975 | none | 1 | 1 | 1 |
| E3 | 34 | 48.4 | 4.0 | 0.0823 | filter | 0.07880 | 0.02864 | 0.01291 |
| E3 | 36 | 33.8 | 4.6 | 0.1353 | filter | 0.26413 | 0.06442 | 0.02570 |
| E3 | 38 | 42.0 | 4.7 | 0.1115 | filter | 0.14718 | 0.04544 | 0.01932 |
| E3 | 40 | 42.9 | 69.3 | 1.6135 | none | 1 | 1 | 1 |
| E3 | 42 | 206.0 | 9.3 | 0.0449 | filter | 0.03218 | 0.01323 | 0.00630 |
| E3 | 44 | 40.1 | 45.5 | 1.1356 | none | 1 | 1 | 1 |
| E3 | 46 | 151.9 | 28.5 | 0.1878 | filter | 1 | 0.14815 | 0.04591 |
| E3 | 48 | 33.1 | 8.3 | 0.2510 | growth | 1 | 1 | 0.09929 |

\* GP14 is an expansion outlier ( $\hat{p}_{\text{obs}} \approx 100$ ) and is excluded from the contraction analysis.
Category abbreviations – filter = filter\_needed, growth = growth\_only\_sufficient, none = no\_contraction, outlier = expansion\_outlier.

#### 4.3.2 Sensitivity to the infection rate

The estimate 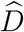 depends on the assumed infection rate *A*, because *A* determines how much extinction is already explained by the growth-only process. Higher *A* increases the growth-only survival probability 1 − *G*(0, Δ*t*), so growth alone accounts for less of the observed diversity loss and more must be attributed to the filter, yielding a smaller (more restrictive) 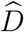. Across the Q1–Q3 range of *A* (1.14 to 2.40 day^−1^), 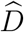 varies by a factor of approximately 3–4 (Table 5). Specifically, the median 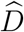 drops from 0.015 to 0.005, and the Q1–Q3 bracket shifts comparably. The Q1–Q3 spread in *A* itself reflects across-animal variation in *λ*, propagated through the *λ* ↦ *A* inversion in Eq. (11). The further amplification to a ∼3–4× spread in 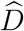 then comes from the inversion in Eq. (19).

Despite this sensitivity, the qualitative conclusion is robust. For the majority of strongly contracting recipients, 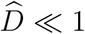 across the full feasible range of *A*, confirming that growth-only dynamics cannot account for the observed diversity collapse without an additional recipient-side bottleneck.

#### 4.3.3 Robustness to the diversity measure

The contraction ratios reported above are the *Shannon-based contraction ratios* 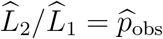, where 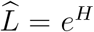 is the *effective number of equally abundant lineages* defined in Eq. (15). For bottleneck analysis this is the biologically meaningful quantity, since it measures how many lineages effectively contribute to the founding population, not merely how many remain detectable. As a sensitivity check, Table 8 also reports the *raw-richness contraction ratio R*_2_/*R*_1_, where richness *R* counts distinct lineages in a sample regardless of how abundant each lineage is. *R*_2_/*R*_1_ is the fraction of lineages that remain detectable from *t*_1_ to *t*_2_. The two ratios measure complementary aspects of contraction. 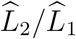 captures whether abundance concentrates into a few dominant lineages, the bottleneck-relevant question, while *R*_2_/*R*_1_ captures whether lineages disappear from detection altogether. Agreement between them, or mechanistically explained disagreement, indicates that the bottleneck conclusion is not caused by the diversity measure we chose. Three observations support this robustness.

First, all six filter-needed E1 animals satisfy *R*_2_/*R*_1_ < 0.12, consistent with 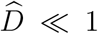 from the Shannon-based analysis. Second, E2’s bimodal structure is preserved. The four contracting animals GP25–28 satisfy *R*_2_/*R*_1_ < 0.75, while the four non-contracting animals GP29–32 satisfy *R*_2_/*R*_1_ ≈ 1.0.

Third, in E3 the two animals where strong Shannon-based contraction is also confirmed by richness are GP42 and GP46, with *R*_2_/*R*_1_ = 0.08 and 0.10. The two measures diverge for four E3 animals, GP34, GP36, GP38, and GP48, where Shannon-based contraction is strong 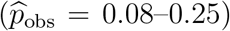 but richness is preserved or grows (*R*_2_/*R*_1_ = 0.85–1.75). This pattern reflects stochastic growth differentiation during early expansion. Random fluctuations cause some lineages to grow much faster than others, so by *t*_2_ a few favored lineages dominate while many others persist at low copy number. As the total population grows, some low-abundance lineages cross the detection threshold, so richness can rise even as Shannon diversity falls. The Hill-based ratio captures this concentration of abundance, the bottleneck-relevant quantity. Raw richness misses it because it weights every detected barcode equally.

Overall, the core finding that E1 recipients require a strong additional filter 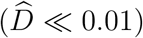 and that E2’s bimodal pattern separates filter-needed from growth-only-sufficient animals is robust to the choice of diversity measure.

## 5 Discussion and Conclusion

The traditional view treats the transmission bottleneck as a sampling step at the moment of physical transfer, in which a small number of virions seeds the new host and sets the diversity of the founding population by sample size alone. This account leaves little room for biological detail in the post-transfer window.

The barcoded influenza A work of Holmes *et al*. [15] refines this picture. They show that high barcode diversity is maintained in inoculated donors and is transferred robustly to recipients exposed by aerosol or direct contact, with many barcodes detectable at the earliest time points positive for infectious virus. Recipient diversity then declines sharply within one to two days after the onset of infection. This temporal pattern places the dominant loss of diversity after physical transfer and during the earliest stages of within-host expansion, rather than at the moment of transmission itself. Host factors active in this window, including innate immune effectors, therefore have greater opportunity to shape the founding population than a pure at-transfer sampling account would allow.

Two quantitative questions follow from this finding. First, is the observed contraction consistent with the intrinsic stochasticity of early viral replication at biologically realistic parameters, or does it require additional loss beyond what growth dynamics alone can produce? Second, to the extent that additional loss is required, how large is it, and how much does it vary between recipients? The size of this residual is the scale of the opportunity available to host factors during the post-transfer window that Holmes *et al*. identify, so answering these questions builds directly on their conclusions. The attachment-lysis branching-process framework of Patwa and Wahl [25], extended here with a recipient-side filter, is what we use to address these questions.

Holmes *et al*.’s temporal repositioning of the bottleneck was qualitative. By making the growth-only extinction probability *G*(0, *t*) computable, our framework shows that what is usually attributed to “few virions reaching the recipient” is actually mostly extinction during the first day of replication. Many virions reach the recipient, but each must survive a eclipse phase under heavy clearance before its first burst, so most fail to produce offspring at all. For roughly half of the recipient animals this growth-stochastic component is sufficient and no additional host biology need be invoked. For the remainder, host biology only needs to account for what remains after growth-only extinction is subtracted.

That residual loss is exactly what the recipient-side filter 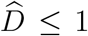 quantifies, on a per-animal basis. This recasts 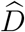 from a phenomenological parameter into an experimental target. As a future direction, pairing barcode-diversity measurements with longitudinal immunological assays would decompose 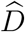 into its immune and non-immune components, turning bottleneck severity into a measurable readout of innate-immune kinetics.

Animal-to-animal variation in transmission outcomes is often attributed to per-host immune differences. Our framework offers a simpler alternative. The 90-fold variation in early growth rate *λ* across recipient animals translates, through an approximately linear composite map (Fig. 2, Right), into a comparable spread in lineage survival probability 1 − *q*. This alone accounts for the two orders of magnitude of variation in observed contraction ratios. Immune heterogeneity may still exist, but it is not required to explain the spread.

The classification of animals into filter-needed and growth-only-sufficient does not depend on the diversity measure used. In Section 4.3.3 we repeat the analysis using raw richness *R*_2_/*R*_1_ in place of Shannon-based Hill numbers, and the qualitative pattern is preserved.

The same framework yields a prediction for vaccinated cohorts. Pre-existing immunity reduces *λ* in each animal, lowering its position on the composite *λ* ↦ 1 − *q* curve. Because 1 − *q* is monotone increasing in *λ*, every animal’s lineage survival drops below the level seen in the naïve cohort. Fewer lineages survive the first one to two days of infection, and the bottleneck is therefore harsher. In a naïve cohort, the few animals with the highest *λ* act as an evolutionary opportunity zone. In these animals, lineage survival is high enough that rare new variants, including potential immune-escape mutants, have a real chance of establishing. Vaccination eliminates this zone. The prediction is directly testable in a vaccinated guinea pig cohort.

Together, these results reframe the transmission bottleneck not as a single sampling event at the moment of transfer but as the cumulative outcome of continuous stochastic dynamics during the earliest phase of infection in the new host. Because the model depends only on four parameters that can be independently estimated for lytic viruses, the approach extends naturally beyond influenza to other systems where transmission bottlenecks constrain viral evolutionary potential.

## A Derivation of the method-of-steps solution for the PGF

Setting *µ* := *A* + *C, δ* := *x* − 1, and *G*(*t*) := *G*(*x, t*), the PGF equation (1) reads

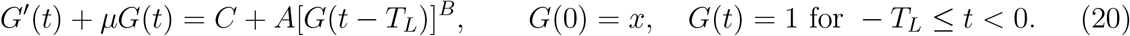

The method of steps [14] solves (20) interval by interval on [*nT*_*L*_, (*n* + 1)*T*_*L*_), *n* = 0, 1, 2, …, because the delayed term depends only on the already known solution on the previous interval. We present Step 0 and Step 1 explicitly, then state the general recursion.

### Step 0 (0 ≤ *t* < *T*_*L*_)

Since *G*(*t* − *T*_*L*_) = 1 on the history, (20) reduces to *G*^′^ + *µG* = *µ*. The integrating factor *e*^*µt*^ with *G*(0) = *x* gives

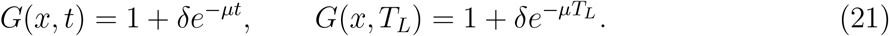

### Step 1 (*T*_*L*_ ≤ *t* < 2*T*_*L*_)

Let *τ* = *t* −*T*_*L*_. Substituting (21) and expanding by the binomial theorem, 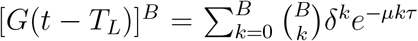. Applying the integrating factor *e*^*µt*^ and integrating from *T*_*L*_ to *t* yields

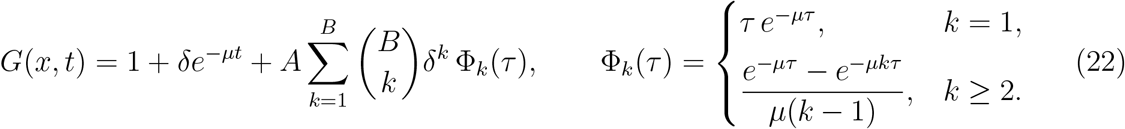

### General step

Let *g*_*n*_(*τ*) := *G*(*x, nT*_*L*_ + *τ*) for 0 ≤ *τ* < *T*_*L*_. Applying the integrating factor *e*^*µτ*^ to (20) on [*nT*_*L*_, (*n* + 1)*T*_*L*_) gives

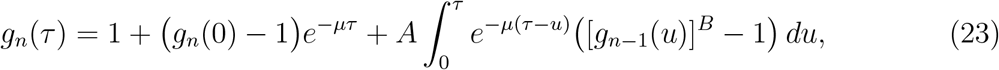

with 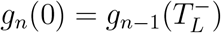. For any 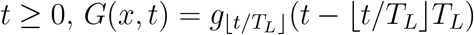.

### Numerical verification

We compared *G*(0, *t*) = *p*_0,grow_(*t*) from the recursion (23) with two independent numerical integrators of (1), namely `solve_ivp` applied interval-by-interval, and the general-purpose DDE solver `JiTCDDE` [2]. The three solutions agree to within 3×10^−7^ over *t* ∈ [0, 1] day (Fig. 4).

## B Supplementary tables

## Acknowledgments

We thank Lindi M. Wahl for her generous support and guidance throughout this work, and for the attachment-lysis branching-process framework [25] that underpins our analysis. We also thank Linda J. S. Allen for her support and for valuable discussions on stochastic processes.

## Code and data availability

All code used to produce the figures and tables in this manuscript is publicly available at https://github.com/wzhan88-1130/Diversity-influenza-bottleneck and archived at Zenodo (DOI: 10.5281/zenodo.20832852). The guinea-pig influenza barcoded-transmission dataset analysed in this work is included as Supplementary Data S2 of Holmes *et al*. [15].

## Notes

https://github.com/wzhan88-1130/Diversity-influenza-bottleneck

https://doi.org/10.5281/zenodo.20832852

### Competing Interest Statement

The authors have declared no competing interest.

